# Cell-body curvature modulates stall frequency to enhance *Vibrio cholerae* swimming and chemotaxis through hydrogels

**DOI:** 10.64898/2026.01.11.698873

**Authors:** Marwan Malik, Ziyang Chen, Andrew G. T. Pyo, Zemer Gitai, Ned S. Wingreen, Marianne Grognot

## Abstract

The swimming motility of *Vibrio cholerae* is a virulence factor that aids in breaching the mucus layer of the small intestine. *V. cholerae* cells have a curved cell shape and previous work demonstrated that loss of curvature decreases infectivity. Here we investigate the mechanism by which *V. cholerae*’s curvature affects single-cell motility within mucus-mimicking environments. Using a multiscale chemotaxis assay, we compared the chemotactic performance of wild-type curved cells (O1 El Tor C6706) and straight mutants under linear chemical gradients in liquid solutions, viscous solutions, and soft agar hydrogels. Our findings reveal that curved and straight *V. cholerae* exhibit similar swimming properties in liquid and purely viscous solutions but significantly differ in hydrogels, with curved cells demonstrating an 86% increase in average chemotactic drift compared to straight mutants in the same chemical gradients. Trajectory analysis indicates that swimming speeds are comparable, but straight mutants experience more frequent stalls, reducing the total time spent swimming. We also found that stalls further reduce chemotactic performances by imposing an average reorientation of bacteria down the chemical gradient, regardless of cell shape. In-silico coarse-grained molecular dynamics simulations corroborate these results and extend them over the wide range of intestinal mucus hydrogel stiffnesses. This model also identifies an optimal curvature for enhanced movement through hydrogel-like meshes that is close to the real median curvature of the pathogen. Our findings thus highlight the mechanisms underpinning cell shape’s role in *V. cholerae*’s pathogenicity and underscore the necessity of studying bacterial behaviors under conditions that simulate host environments.

## Introduction

*Vibrio cholerae*, the causative agent of cholera, remains a global human health concern. This bacterium has a characteristic curved (“comma-shaped”) morphology and can swim by rotating a single polar flagellum. Its swimming motility has been recognized as a virulence factor^1,2^, likely for allowing *V. cholerae* to swim through the mucus protecting the small intestine^3^ during the early steps of infection. Two genes (*crvA* and *crvB*) have been shown to directly determine *V. cholerae*’s body curvature^4^. Their deletion brings its cell-body shape to a straight rod. A straight-shaped *crvA* mutant displayed an infectivity defect in two animal models of infection^5^.

The effect of *V. cholerae* curvature on infection could potentially be explained by the curved shape enhancing swimming performance in hydrogels such as mucus to improve its penetration rate and therefore increase pathogenicity. In simple liquid medium, one study demonstrated a marginal (approx. 5%) increase in swimming speed due to cell-body curvature^6^. However, the effect of *V. cholerae* cell shape on single-cell swimming behaviors has not been studied previously in hydrogel environments more closely mimicking the host environment. This gap is likely due to the difficulty of collecting the large number of long single-cell trajectories necessary to resolve navigation performances, especially in complex media. Navigation performances are understood here as encompassing both swimming motility (random movement) and chemotaxis (biased movement in chemical gradients).

To meet this challenge, we made use of a recent multiscale chemotaxis assay^7,8^. In this assay, single-cell behaviors are 3D tracked under a well-defined linear chemical gradient in liquid solutions, viscous media, or agar hydrogels. A dilute agar hydrogel was used as the main mucus-mimicking environment, owing to its hydrogel structure with pores of similar size to pores in intestinal mucus hydrogels (see SI Discussion). The assay simultaneously resolves population-scale performances (e.g. chemotactic drift speed) and how individual cells swim in order to achieve such performances. By comparing the curved phenotype (wt strain) against the straight-shaped phenotype (Δ*crvAB* deletion mutant strain), we aimed to directly resolve the potential boost offered by the curved shape compared to the straight rod shape, and pinpoint which navigation properties (speed, turning rate, turning angle, stop duration, etc.) are leveraged to do so. We also developed an in-silico coarse-grained molecular dynamics model of curved or straight cells moving and rotating through a hydrogel-like mesh. By varying cell curvature and properties of the mesh, we were able to dissect the biophysical parameters that govern how effectively cells traverse crowded or obstructive surroundings, thereby complementing and extending insights gained from our experimental observations.

## Results

### Chemotactic performances in buffer are independent of cell-body curvature

Wild-type (wt, O1 El Tor C6706) and straight Δ*crvAB* strains were grown to late exponential phase, to maximise curvature of wt^4^ (**Figure 1a**). High bacterial densities (yielding high curvature) correlated with a high motile fraction and fast swimming speed (SI Figure 1a). Overall, in buffer environments, wt and Δ*crvAB* strains exhibited remarkably similar swimming properties with indistinguishable chemotactic performances in a 200 µM/mm L-serine gradient.

**Figure 1.**
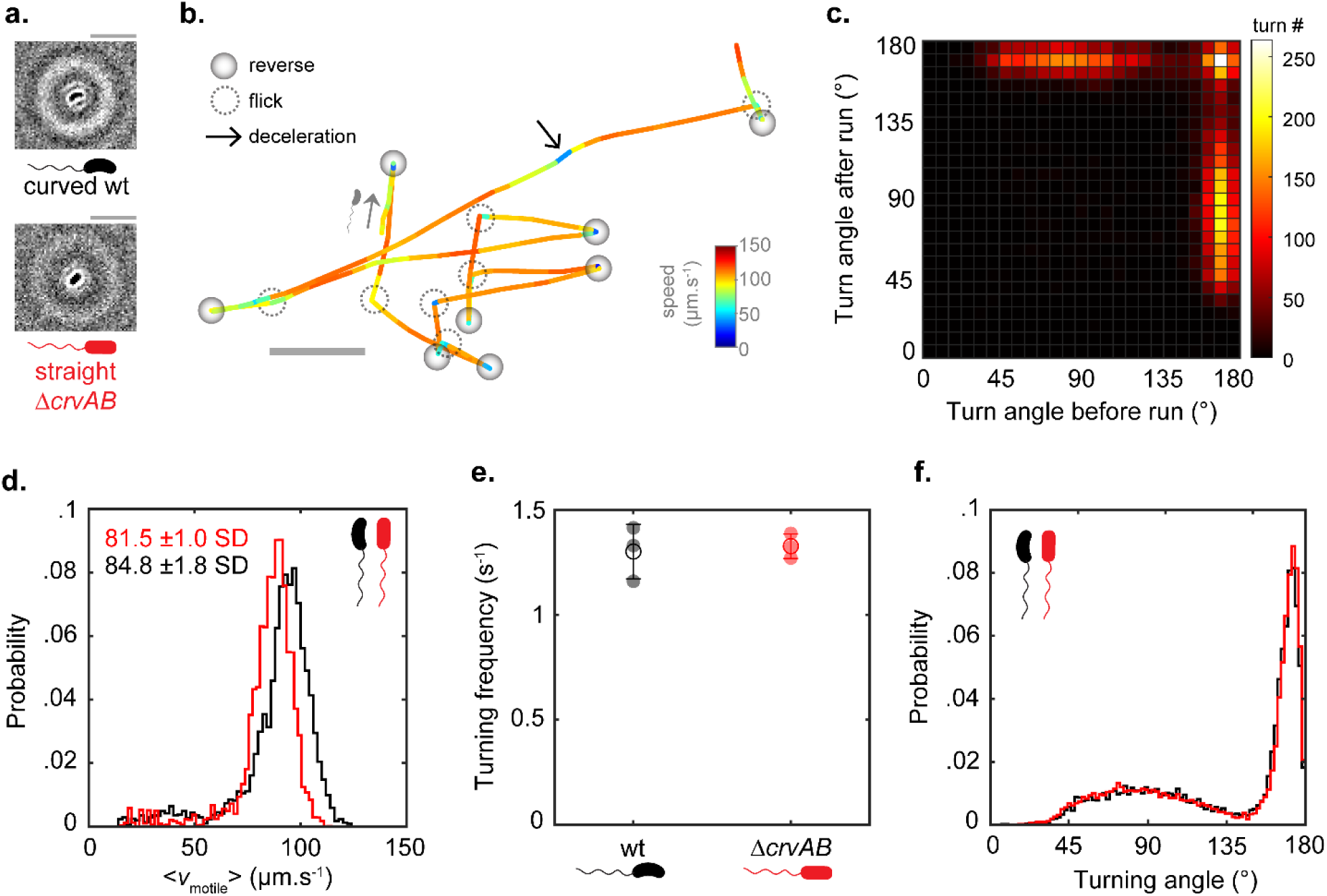
In buffer, *V. cholerae* C6706 str2 wild-type (wt) and its non-curved mutant Δ*crvAB* show run-reverse-flick motility with similar properties. **(a)** Phase-contrast images from 3D-tracking acquisitions (see Methods) of the two phenotypes, and the associated symbols subsequently used. Scale bars represent 8 µm. **(b)** Example 6.66-s trajectory of wt strain from 3D tracking in motility medium (M9TB), with color reflecting instantaneous speed. Turning events identified as reverses (above 150°) or flicks (under 140°) are indicated by closed circles and dotted circles respectively, while a deceleration event during a run is indicated by a *black* arrow. The start of the trajectory is indicated by a grey arrow and a forward-swimming bacterium. **(c)** Example bivariate histogram of consecutive turning angles observed after versus before the same run, for the wt strain. The distribution confirms run-reverse-flick motility, where reversals can be preceded and followed by reversals or flicks, but two flicks never occur in a row. **(d)** Average swimming speed distributions of motile wt (black) and Δ*crvAB* (red) with averages and standard deviations over three biological replicates. Turning frequencies **(e)** and turning angle distribution **(f)** of wt (black) and Δ*crvAB* (red) are indistinguishable. The full characteristics of the above-analyzed biological triplicates are available in *Error! Reference source not found.*.

We first analyzed random single-cell motility by high-throughput 3D tracking in homogeneous liquid motility medium lacking a chemotactic gradient (M9 salts with 0,01% TB, M9TB) over three biological replicates (detailed datasets available in SI Table 1). These experimental conditions were chosen to ensure stable motility properties over at least 130 minutes (SI Figure 1) within which all later acquisitions were made. Wt and Δ*crvAB* displayed similar average swimming speed within a few micrometres per second (**Figure 1d**). We applied an automated turn event detection to trajectories of at least 0.8 s and with speed over 15 µm/s (see Methods) to further analyse the random swimming properties. Both strains displayed a run-reverse-flick motility, where turning angles are characterized by a succession of reversal and flick or reversal and reversal but never two flicks in a row (**Figure 1b,c**). Decelerations were occasionally visible in trajectories during runs (**Figure 1b**, arrow) as previously described for *V. cholerae* strain O395^9^. Turning frequencies and turning angle distributions were indistinguishable between the curved and the straight cell-body phenotypes (**Figure 1e,f**).

Chemotaxis to L-serine was captured by a multiscale chemotaxis assay: bacteria were 3D tracked in the middle of a 200 µM/mm linear serine gradient established by pure diffusion between two reservoirs containing dilute bacteria with or without 200 µM serine in M9TB medium (**Figure 2a**). From the thousands of trajectories, we found that the average swimming speed of wt is 4.2(0.6 95% CI)% higher than that of Δ*crvAB* (SI Figure 2a-b), similar to previous observations by Fernandez et al.^6^. However, this small difference did not result in any significant chemotactic advantage for the curved-cell shape over its straight-cell counterpart (**Figure 2b,c**). Specifically, the populations’ chemotactic performances, defined here as their chemotactic drift velocity (average population’s displacement up the gradient), showed no significant difference between wt and Δ*crvAB* (**Figure 2b,c**). On average, the chemotactic drift of the straight-shaped population was 103(±7 95%C.I.)% that of the curved one over ten replicates.

**Figure 2.**
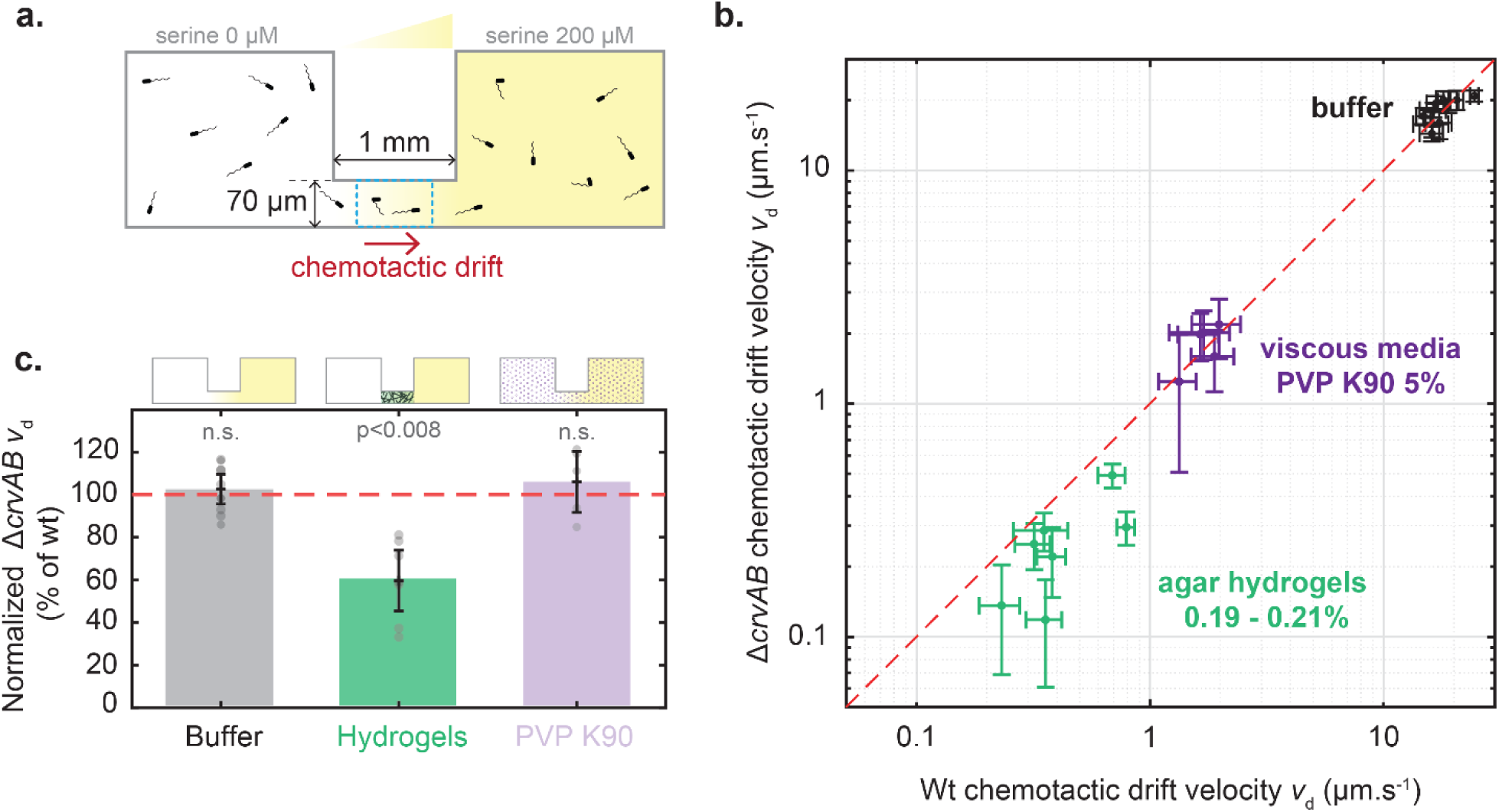
Population-scale chemotactic performances in buffer and mucus-mimicking media. **(a)** Schematic of the multiscale chemotaxis assay (see also Grognot et al.^7,8^). Bacteria are 3D tracked in the middle of a microfluidic channel (blue box) where a linear serine gradient is established by diffusion between two reservoirs with equal bacterial concentrations, but with or without 200 µM L-serine. The chemotactic drift of the population is its net speed up the 200 µM/mm serine gradient, computed as the average of all signed instantaneous velocity components along the gradient direction. **(b)** Comparison of chemotactic drift velocity of curved wt against non-curved Δ*crvAB* mutant in buffer (black, n=10), viscous polymer PVP K90 (purple, approx. 38 centipoises, n=3), or soft agar hydrogels (green, n=7, 0.19-0.21% (w/v)). Each point is an independent replicate, representing a pair of drift measurements for both phenotypes, acquired on the same day in the same media. Error bars reflect 95% confidence intervals estimated by a jackknife resampling procedure consisting of dividing the data into subsets of 200 trajectories and computing the SEM drift obtained for different subsets. The dotted red reference line marks identical performance of both phenotypes. **(c)** Normalized chemotactic drift velocity of Δ*crvAB* relative to wt from the same biological replicate, in buffer (grey, n=10), PVP K90 (purple, n=3), or agar hydrogels (green, n=7). Indicated p-values are from a paired *t*-test with unknown variance. Datasets used in panels b and c are detailed in SI Table 2.

### Cell curvature enhances chemotactic performances in soft hydrogels

In contrast to liquid environments, experiments conducted under the same 200 mM/mm L-serine gradient but within the structured environments of agar hydrogels (0.19-0.21% w/v) revealed notable differences between the chemotactic performances of the wt and Δ*crvAB* strains. The curved phenotype outcompeted its straight-shaped counterpart: out of seven experimental replicates, only one instance showed no significant difference in chemotactic drift within 95% confidence intervals (**Figure 2b**). On average, the chemotactic drift of the straight-shaped population was only 60(±7 95% CI)% that of the curved one (**Figure 2c**).

We also compared chemotactic performances in highly-viscous polymer solutions of PVP K90, a simple linear polymer with non-Newtonian behaviors^10^, at bulk viscosity of approximately 38 cP (5% w/v, see Methods). No significant difference in chemotactic drift was observed between the two phenotypes over 5 experimental replicates, with on average a chemotactic drift of the straight-shaped population 106(±14 95% CI)% that of the curved one. Trajectory inspection revealed a slow run-reverse-flick motility, as reported previously in other strains^9^. The average swimming speeds were also similar, at 11.9 µm/s and 12.0 µm/s for the wt and Δ*crvAB* strains respectively. When further comparing average speeds or chemotactic performances for subpopulations of similar speeds, no significant differences were found (SI Figure 2c,d). We conclude that the presence of uncrosslinked polymers per se does not lead to a chemotactic advantage for curved cells. Because the mesh structure of hydrogel seems to be important for curved cells to outperform their straight counterparts we focused the rest of our work on the analysis of navigation in hydrogels to investigate the mechanism(s) behind the clear outperformance by the curved cell body.

### Cell curvature improves performances in soft hydrogels by increasing swimming extents

Analysis of single-cell trajectories in agar hydrogels revealed a distinct swim-stall motility pattern (**Figure 3a**), similar to another polarly-flagellated Vibrio, *Vibrio alginolyticus*^8^. To distinguish swimming phases from stalling phases we ran an automated trajectory analysis (see Methods) on each of the many trajectories yielded per experimental replicate (see detailed datasets SI Table 2). Phases were defined as periods of 0.2 s or more at instantaneous speed over (swimming) or under (stalling) 1 µm/ s (**Figure 3b**). Notably, swimming speeds and root-mean-squared swimming speeds were not significantly different between wt and Δ*crvAB* strains (**Figure 3c**). In contrast, there was a marked increase in stall frequency for straight bacteria (**Figure 3d**). This result indicates that cell-body curvature enables *V. cholerae* to swim significantly further before encountering a stalling event (**Figure 3e**). Additionally, despite similar instantaneous swimming speed, the chemotactic speed during swimming phases was significantly lower for the straight mutant (**Figure 3f**). The overall decrease in chemotactic performance in the absence of curvature (by approx. 40%) can be understood as a result of two factors: an increase by ∼10% of time not moving, and a decrease by ∼35% of chemotactic efficiency during swimming. In other words, decreasing swimming extents not only increases the time spent stuck in a stall, but leads to worse performance between stalls.

**Figure 3.**
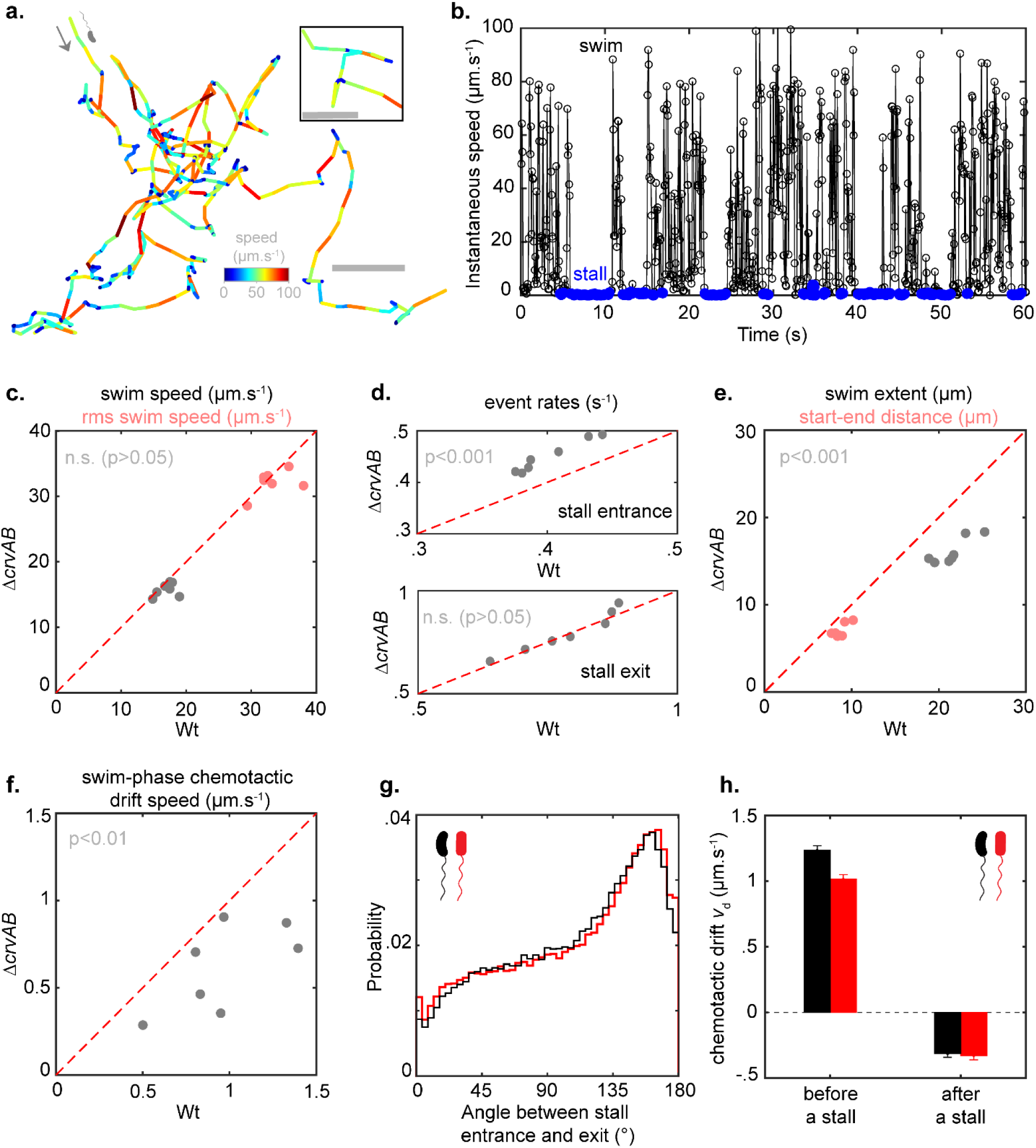
Swim-stall motility in soft agar. **(a)** A 1-min long example trajectory for the curved phenotype (wt) with an average speed of 19.5 µm/s. Inset shows a segment of trajectory with run-reverse-flick appearance. **(b)** Time trace of instantaneous speed for the same trajectory. Swim (black) and stall phases (blue) begin when the bacterium’s instantaneous speed is above, respectively below, 1 µm/s for at least 0.2 seconds. **(c-e)** Swim-stall motility parameters averaged over each independent replicate and each phenotype: rms swim speeds (c), stall frequency over swim phases (d), and swim extent computed either as cumulative distance travelled during a swim phase (e, black) or as the physical distance between beginning and end of a swim phase (e, pink). **(f)** Average chemotactic drifts computed over the swimming phases only. Indicated p-values are from paired *t*-tests with unknown variance. The dotted red reference lines mark identical performance of both phenotypes. **(g)** Angle between the entrance direction and the exit direction of stall phases, computed respectively on two instantaneous speed vectors before and two after the stall. The median angle was 122° and 126° for wt and Δ*crvAB* respectively. (**h**) Chemotactic drift velocities just before or just after a stall, respectively from two instantaneous speed vectors before and two after the stall, for wt (black) or Δ*crvAB* (red) strains, averaged over all replicates in agar hydrogels. Error bars indicate the standard error of the mean. Analysis included all fully defined (i.e. with a start and an end) stalls detected over the n=7 replicates, totaling 103,677 and 95,151 stalls for wt and Δ*crvAB* respectively.

### Cell curvature helps avoid stalls but turning rates determine escape from stalls

In our analysis, no significant effect of cell-body curvature was observed on stall durations, with an average of 1.28 (±0.16 SD) and 1.27 (±0.16 SD) seconds for curved and straight shapes, respectively. These observations prompted us to interrogate further the mechanisms of stall escape. We hypothesized that once bacteria enter a stall state, turning (i.e. reversing or flicking) dominates the chance of successful escape. Turning rates were similar for curved and straight bacteria in buffer, which would explain their similar stall escape rates in hydrogel. Consistent with this hypothesis, the escape rate of stalls (approx. 0.8 s^-1^) is close but lower than the turning rate observed in buffer (approx. 1.3 s^-1^). Similarly, the distribution of angles at which a bacterium escapes a stall (relative to its direction upon entering it) is reminiscent of the typical distribution of reverses and flicks in buffer (**Figure 3g**) as also qualitatively observed in **Figure 3a** (inset).

To further test our hypothesis, we induced a reduction in turning frequency by imposing a serine shock treatment (50 mM) to wt bacteria in an agar hydrogel. In buffer, the sudden presence of 50 mM serine in motility medium sharply decreased turning frequency, especially during the first 5-10 minutes, and for at least 30 minutes (SI Figure 4a). Similar imperfect adaptation to high concentrations of serine was previously observed in *Escherichia coli*^11^. In agar hydrogel, the sudden presence of 50 mM serine in both reservoirs led to a 1.5-fold increase in average stall durations (SI Figure 4b) relative to before shock. These findings suggest that in hydrogels cell-body curvature facilitates avoidance of stalls while cell curvature-independent turning is the main mechanism for escaping stalls.

### Stalls decrease chemotactic performance by reorienting bacteria down the chemical gradient

We next sought to understand our observation that stalls not only reduce time spent moving but also decrease chemotactic efficiency during swimming phases (**Figure 3f**). We hypothesized that the effect of stalling on chemotaxis could be related to the turning-based mechanism of stall escape. The average chemotactic drift of bacteria just before entering a stall is positive, while it is negative (and similar between curved and straight strains) at the exit of a stall (**Figure 3h**). As such, a stall is not a simple reset of chemotactic sensing randomly reorienting the bacterium, but an adverse event that on average tends to orient bacteria away from chemotactic gradient. This can be understood because bacteria are typically oriented towards the gradient when they stall, but to escape the stall they require a turn that typically changes their angle more than 90° away from their initial trajectory (**Figure 3g**). In other words, a bacterium is more likely to be swimming up the gradient before a stall, which means that bacteria initially escaping a stall are more likely to be moving in the “wrong” direction (down the gradient). Consequently, the ability of cell curvature to reduce the rate of stalling increases chemotactic performance by both increasing the time spent swimming and decreasing the encounter of events negatively affecting the performances of the following swimming phase.

### An in-silico model reveals a biophysical mechanism for curvature-enhanced motility in hydrogels and highlights a curvature optimum

To further interrogate the biophysical mechanism(s) underpinning our findings, we developed an in-silico coarse-grained molecular dynamics model of curved or straight cells moving and rotating through a hydrogel-like mesh (see Methods). To simplify the model and focus on the key parameter of stall onset frequency, reversals and flicks were not included: once the forward-swimming model “bacterium” enters a stall, it never escapes. The in-silico hydrogel was characterized by an average pore size of 3.49 μm (±1.79 μm SD, SI Figure 6a), within the estimated range of mean pore size for soft (0.25%) agarose hydrogels^12^ and close to recent estimates of median pore sizes of canine intestinal mucus using Cryo-SEM^13^. We kept these pore sizes constant but modified the elasticity of the links between mesh elements to screen a larger range of model hydrogel stiffnesses, covering the expected range of soft agar hydrogels (hundreds of Pa) but also the expected typical 1-100 Pa stiffness range of human intestine mucus.

When comparing a thousand trajectories from straight or curved (0.71 µm^-1^) “bacteria”, the model replicated the experimentally observed boost in travel distance before stall conferred by curvature. Upon estimation of exponential decay lengths of these travel-distance distributions (dashed lines in **Figure 4a**, summary in SI Table 3), the model returns an exponential decay length ratio of 1.25, which despite the model’s simplicity is in good quantitative agreement with experimental values of 1.26 (±0.10 SD, n=7). This curved-cell advantage is robust over a wide range of mesh stiffnesses (**Figure 4b**). As stiffness decreases, the model even predicts a higher advantage for the curved shape, with a ratio increasing up to almost 1.5 for shear modulus close to human small-intestine mucus (under 10 Pa).

**Figure 4.**
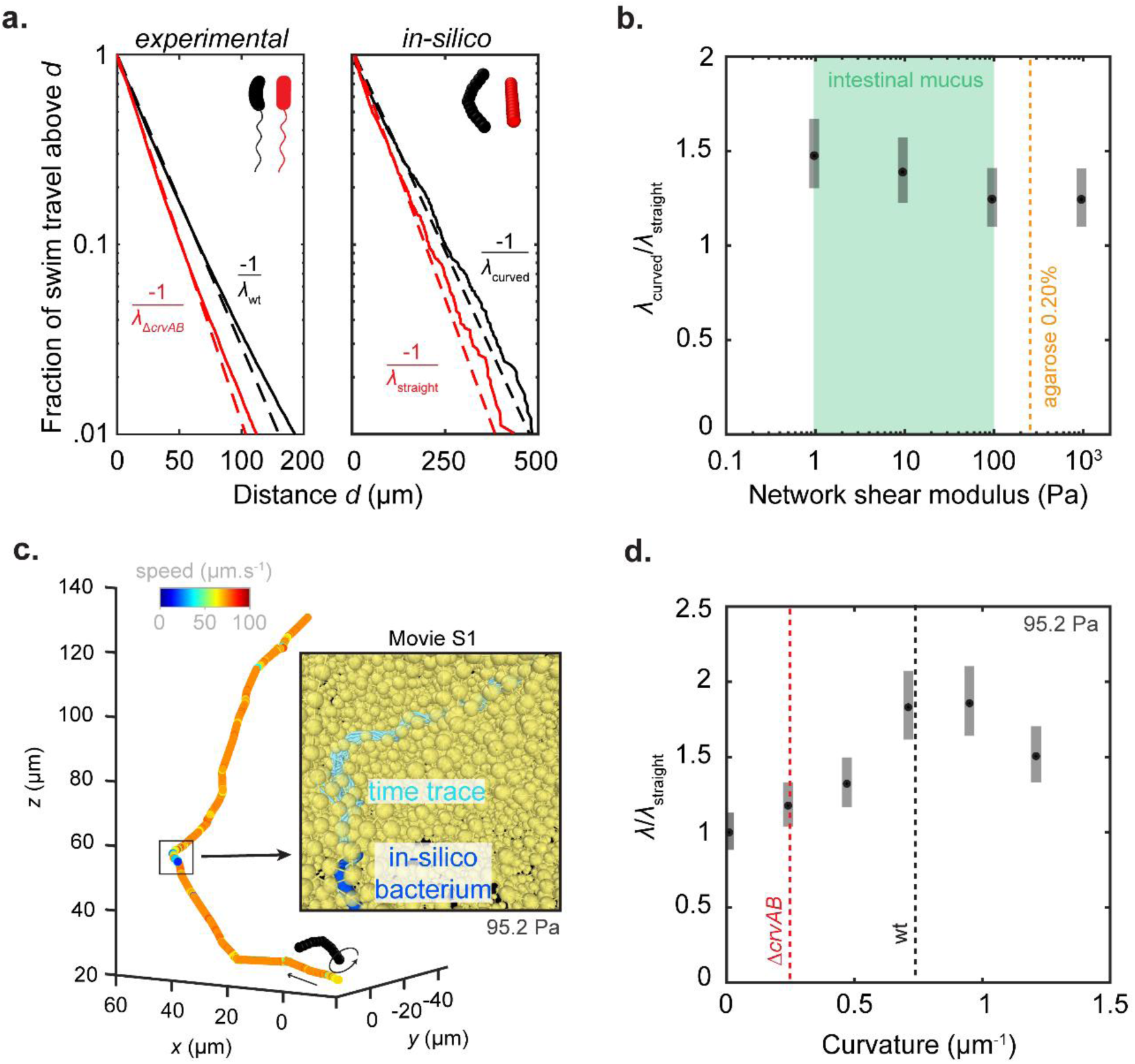
Curved shape advantage in in-silico model. **(a)** Survival distributions of travel extents in experimental data (left) or in a coarse-grained molecular dynamics model (right, n=1000 simulations for each cell shape). Dashed lines represent the slopes from the estimates of exponential decay lengths *λ*. In the in-silico model, curvatures are 0 and 0.71 µm^-1^ for curved and straight cells respectively, and the network’s shear modulus is 95.2 Pa. **(b)** Ratio of decay lengths *λ* for curved (0.71 µm^-1^) relative to straight in-silico bacteria in networks of different shear modulus obtained by modification of the properties of the mesh (the higher the deformability, the lower the shear modulus). Error bars represent the extrema of combined 95% confidence intervals determined for each *λ* estimate used in the ratio. The orange dotted line denotes the shear modulus of agarose 0.20% reported by Shahab et al.^14^. **(c)** Representative example trajectory from one simulation with curved shape, including a slower phase that doesn’t result in a stall. **(c inset)** Detail of the trajectory segment with strongly decreased speed, where the in-silico bacterium (dark blue) can be seen rotating (light blue segment for each bead) and avoiding a complete stall within the network (yellow). The corresponding movie is provided to clarify further the observed effect, see SI Movie 1. **(d)** Ratio of decay lengths *λ* for in-silico bacteria with different curvatures, relative to *λ*_straight_. A ratio significantly over 1 signifies that the straight shape is outperformed. The black dotted line denotes the experimental median curvature reported for wt in late exponential phase by Martin et al.^4^. Error bars represent the extrema of combined 95% confidence intervals determined for each *λ* estimate used in the ratio.

Importantly, the model allowed us to precisely visualize cell-body movement in the mesh, as opposed to our experimental techniques. We observed multiple events of curved cells decelerating, sometimes to an almost complete stop, followed by a screw-like process that leads to a direction change and a return to previous speed without stalling (see example in **Figure 4d** and SI Movie 1). While both straight and curved cells can slide along a single obstacle, straight cells may get stuck in pocket-like traps formed by multiple obstacles, whereas curved cells can often adjust their attack angle to avoid a stall. We conclude that the advantage of model curved cells lies in their ability to avoid true stalls by deflecting their trajectory away from the trap, preventing themselves from stalling.

Interestingly, upon variation of cell curvature, the model also highlighted an optimum in cell curvature to avoid stalling (**Figure 4d**). This optimum is close to realistic 3D curvature values for the wt strain in late exponential phase published by Martin et al.^4^ with a median of 0.75 µm^-1^. At higher curvature, the advantage decreases, which suggests that cells with more U-shaped geometries do not efficiently reorient themselves to escape traps.

## Discussion

Our work highlights the significant role of cell-body curvature in enhancing the chemotactic performance of *V. cholerae* within hydrogels, as well as the underlying mechanisms at play. Its curved cell body allows *V. cholerae* to extend swimming phases between stalls when navigating hydrogel meshes under a chemical gradient, thereby improving its ability to cross the hydrogel (by approx. 40% here). Altogether, our work extends the understanding of how curvature improves motility and chemotaxis by addressing these behaviors in more complex and realistic environments. We suggest that our findings are not only relevant to understanding how *V. cholerae* crosses the mucus hydrogel protecting the human small intestine but also for the pathogen’s escape from biofilms, as biofilms also represent hydrogels whose pore sizes can be in the range of those hindering bacterial movements^15^.

Our computational model allows to extend the role of curvature in boosting navigation over a wider range of hydrogels, beyond experimental reach. It also provides some mechanistic insight into how curved cells avoid traps. Notably, the naïve intuition that a curved cell acts as a “screw” boring through the simulated hydrogel is only partially borne out. Unlike the motion of a screw, the rotating curved cell can change its angle of attack to avoid being trapped. This is evident in the example in **Figure *4*d** (see also Movie S1). The model also reveals an optimal cell curvature close to that observed in real *V. cholerae* cells. Within the model, increasing curvature beyond the optimum has two drawbacks: First, increasing curvature slows the average speed of cells through the hydrogel (SI Figure 5a), which is expected from the extra collisions of the rotating cell with the hydrogel. However, this is only a small effect (∼10%) over the range of curvatures considered. Instead, the more highly curved cells fail to find alternative paths through the simulated hydrogel. Since the model is only a rough simulacrum of a real hydrogel, how high curvature affects escape from traps in real systems, including mucus or *V. cholerae* biofilms, remains a topic for further exploration.

Our findings draw an intriguing parallel with *Vibrio alginolyticus*, which employs a different strategy to achieve similar enhancements in swim extents despite lacking cell-body curvature. Instead, *V. alginolyticus* features a dual flagellar system that includes both polar and lateral flagella. The presence of lateral flagella has been shown to increase swimming distances in soft agar hydrogels (0.15-0.25%), resulting in better chemotactic performance compared to a population expressing only a polar flagellum^8^. Interestingly, while both body-curvature and lateral flagella improve swim extents in hydrogels, lateral flagella also improve swimming and chemotaxis of *V. alginolyticus* in high-viscosity PVP K90 while cell-body curvature of *V. cholerae* does not. Another notable difference is that *V. cholerae*’s curvature does not hinder navigation in buffer (compared to straight shape), while *V. alginolyticus*’s lateral flagella significantly decrease swimming speed and chemotactic performances (compared to the polar-flagellum-only phenotype). Expression of lateral flagella also entails significantly reduced growth rates, while curvature does not. In the *Vibrio* family, cell-body curvature and dual flagellation appear to be mutually exclusive: the three known Vibrio species lacking CrvA/CrvB homologs^4,5^ have dual flagellar architectures^16–18^ while none of the species containing CrvA/B homologs are known to have lateral flagella. Might body curvature and lateral flagella, while both efficient ways to improve hydrogel penetration, be incompatible, or are they two independent strategies to improve navigation properties under different selection criteria? For example, curvature could represent a way to specifically improve infectivity without significantly reducing growth or becoming more immunogenic by increasing flagellin number, while lateral flagella could promote motility in more diverse environments at a higher adaptative cost. Altogether, these observations suggest a fascinating and complex landscape of bacterial adaptation where cell-body curvature and lateral flagella are two distinct exemplary strategies for improving bacterial navigation. Our work ultimately highlights the importance of interrogating navigational properties in host-relevant environments, here by combining a high-throughput experimental method with parallel mechanistic simulations.

## MATERIALS & METHODS

### Bacterial strains and growth conditions

*V. cholerae* C6706str2 wild-type (wt) and its non-curved mutant obtained by point mutations in curvature-inducing complex proteins CrvA and CrvB (Δ*crvAB*) originated from Martin et al.^4^. Overnight cultures were inoculated in 2 ml LB (1% Bacto Tryptone, 0.5% Bacto Yeast extract, 0.5% NaCl, pH 7) from individual colonies grown on 2% agar LB plates streaked from frozen glycerol stocks less than 1 week earlier, and grown to saturation at 30°C, 250 rpm. Day cultures were inoculated with the overnight cultures at 1:200 dilution in 10 ml TB (1% Bacto Tryptone, 0.5% NaCl, pH 7) in and grown in parallel at 30°C, 250 rpm in 100-ml glass culture flasks with ventilated caps (DWK Life Sciences), until they reached an optical density (OD) at 600 nm between 2.2 and 2.4 and within 0.01 of each other (to ensure comparability, see SI Figure 3). Volumes of 0.5 ml of bacterial culture mixed with 0.5 ml of M9TB were washed once by centrifugation in 1.5 ml microcentrifuge tubes (7 min at 2,500 rcf), followed by gentle resuspension in 1 ml of M9TB. M9TB consisted of M9 salts (M6030 Sigma-Aldrich) sterile filtered and 0.01% (v/v) TB (autoclaved) added to it. Bacterial suspensions were diluted to a target OD of approximately 0.005 for chemotaxis experiments in buffer, 0.003 for polymer solutions, or 0.002 for chemotaxis experiments in agar, either in M9TB (chemotaxis experiment in buffer or agar) or in M9TB with PVP K90 (chemotaxis experiment in viscous media), with or without chemoattractant, for injection into the chemotaxis device. For bulk viscosity estimates of 5% PVP K90 solutions, a previously established relationship was used^9^.

### Chemotaxis assay preparation in buffer or polymer solution

The chemotaxis assays were performed using a high-throughput chemotaxis assay^7^ where bacteria are tracked in 3D in a controlled chemical gradient established in a commercially available microfluidic device (µ-slide Chemotaxis, IBIDI), consisting of two reservoirs of different chemical concentrations connected by a small channel in which a linear gradient is formed. The device’s reservoirs were respectively filled with bacterial solutions with or without 200 µM L-serine (84959, Sigma-Aldrich), as previously described^8^, so as to obtain a 200 µM/mm stable and repeatable linear gradient within approximately 40 minutes after closing the device^7^.

### Chemotaxis assay preparation in agar

For chemotaxis assays performed in agar, the central channel of the microfluidic device was filled in advance with molten agar (0.19-0.21% w/v Bacto agar in M9TB). For filling the device, each reservoir is successively emptied, filled with bacterial solution with or without chemoattractant, and then closed. Biological replicates comparing phenotypes at a given agar concentration were prepared in parallel to limit the influence of technical variability in agar gel properties. Chemotaxis experiments in buffer and polymer solutions were concluded typically within 60 min after closing the chamber for experiments. Experiments in agar concluded typically within 1.5h, maximally 2h.

### Motility experiments

After washing and resuspension in M9TB as described above, cells were diluted to a target OD of approximately 0.005 in M9TB, incubated at room temperature for at least 15 minutes, to allow adaptation to the medium. Preliminary experiments followed similar protocol but with different resuspension media or preincubation time as indicated in Supplementary Figure 1. Sample chambers with a height of approximately 300 µm were created as described previously^31^, by using three layers of parafilm as spacers between a slide and a cover glass, filled, sealed with valap and immediately brought to the microscope for data acquisition.

### Data acquisition

Phase contrast microscopy recordings were obtained at room temperature (∼21°C) on a Nikon Ti2-E inverted microscope using an sCMOS camera (PCO Edge 5.5) and a 40x objective lens (Nikon CFI S Plan Fluor ELWD 40x ADM Ph2, correction collar set to 1.2 mm to induce spherical aberrations^31^) focused at the center of the channel in all three dimensions. The integration time was 3 ms. The frame rate was at 30 fps for recordings in buffer, 15 fps in agar and 5% PVP K90 solutions. Typical cumulated acquisition time for one condition in one chemotaxis experiment (one point in Figure 2b,c) ranged from 4 to 10 minutes in buffer, 6 to 12 minutes in viscous medium, 15 to 20 minutes in agar. All acquisitions were made within 150 minutes of resuspension in motility medium after washing.

### Data processing and 3D tracking

Video recordings were binned by a factor of 2 × 2 pixels by averaging pixel counts and then subjected to a background correction procedure based on dividing the image by a pixel-wise median computed across a sliding time window of 101 frames, except for data acquired for agar experiments, where a sliding window of 2001 frames was used to ensure that stalled bacteria were not erased. 3D trajectories were extracted from phase-contrast recordings using a high-throughput 3D tracking method based on image similarity between diffraction rings of bacteria and those in a reference library with known vertical positions^31^ improved in previous works by correcting images for the diffraction rings of already tracked bacteria^8^.

Positions were smoothed using 2^nd^ order ADMM-based trend filtering^32^ with regularization parameter *λ* = 0.1 in agar and *λ* = 0.3 otherwise. Three-dimensional velocities were computed as forward differences in positions divided by the time interval between frames. In chemotaxis experiments, the vertical position of the top and bottom of the chamber were identified by visual inspection of trajectory data. All trajectory segments within at least 12 µm of the top or bottom of the central channel were removed to avoid surface interaction effects^7^.

### Run-reverse-flick analysis in buffer

The turning event detection, that we previously applied to another *V. cholerae* strain (including sharing of the code)^9^, is based on the local rate of angular change, computed from the dot product between the sums of the three consecutive velocity vectors preceding and subsequent to a time point. The threshold for a turn to begin is a 7-fold rate relative to the median rate of angular change of the run segments, as determined in three iterations of the procedure. A new run begins with at least one time points (at least 0.033 s) under this threshold. The 3D turning angle for a turn beginning at frame *i* and ending at frame *j* is computed as the angle between the sum of the instantaneous velocity vectors at frames *i*-2 and *i*-1 and the sum of those at frames *j* and *j*+1. Backward (CCW rotation) and forward (CW rotation) runs were identified as runs with a turn under 140°, respectively at the end or at the beginning of the run, and a turn above 150° at the other end of the run. All trajectories with a minimum speed of 15 µm/s and with a minimum duration of 0.8 s were analyzed.

### Stall-swim analysis in soft agar hydrogels

The stall event detection is based on the instantaneous velocity: If the speed is under 1 µm/s for 0.2 s or more, a stall starts. During a stalling event, if the instantaneous speed exceeds 1 µm/s for 0.2 s or more, the stall ends, and a new swimming phase starts. All trajectories with a minimum duration of 1 s are analyzed.

### Simulation model

Coarse-grained molecular dynamics simulations are conducted using the Large-scale Atomic/Molecular Massively Parallel Simulator (LAMMPS^33^). In the simulations, the gel matrix is represented as a crosslinked network of chains of spherical beads of 1 µm diameter and 9.42478 pg mass. The beads within a polymer chain are interconnected by stretchable bonds, defined by energy:

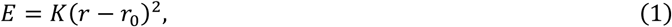

where *K* is 2.2 nN/µm and *r*_0_ is 1.25 µm. The swimming cell is modeled as a rigid body consisting of 13 spherical beads each of 1 µm diameter and 1 pg mass. For the straight cell, all beads are aligned linearly, whereas for the curved cell, the beads form an arc. Interactions between gel-cell beads and gel-gel beads are modeled by a purely repulsive Lennard-Jones potential defined as:

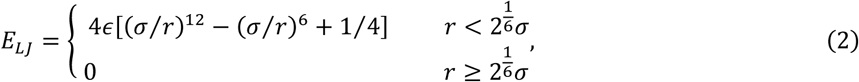

where *ɛ* is 0.0001fJ, *σ* is 1 µm. The equations of motion are integrated in the microcanonical ensemble (NVE), ensuring conservation of energy through direct integration of Newton’s equations. Simultaneously, a Langevin thermostat is applied to maintain the system at each desired temperature (see below).

### Gel initialization

First, a gel matrix formation process is simulated. 125 chains each consisting of 150 beads are introduced into a simulation box of dimension 100 µm×100 µm×100 µm with periodic boundary conditions. To better replicate the compactness and pore size distribution typical of a hydrogel, the system is geometrically transformed: all bead coordinates (*x*, *y*, *z*) and the simulation box are scaled by a factor of 0.5 to uniformly shrink the system, generating a compact initial configuration. The system is then fully relaxed for 3×10^5^ time steps (the timestep is 1 µs) at 300 K. Following this, crosslinking between gel beads is allowed during 3×10^5^ time steps, while the system is cooled down from 300 to 100 K. During the crosslinking phase of the simulation, bond formation is implemented by the *fix bond/create* command in LAMMPS. A check for possible new bonds is performed every 100 time steps. If two beads *i* and *j* are within a distance of 1.125 µm of each other, and if a bond does not already exist between *i* and *j*, and if both *i* and *j* are below their respective maximum bond limit (no more than 4 bonds), then *i* and *j* are labeled as a “possible” bond pair. If several beads are close to a bead, it may have multiple possible bond partners. Every bead checks its list of possible bond partners and labels the closest such partner as its “sole” possible bond partner. After this is done, if bead *i* has bead *j* as its sole possible partner and bead *j* has bead *i* as its sole possible partner, then the *i*,*j* bond is formed with a probability of 0.5. Note that these rules mean that a bead will only be part of at most one newly created bond at a given time step. It also means that if *i* chooses *j* as its sole possible partner, but *j* chooses *k* as its sole possible partner (because distance*_jk_* < distance*_ji_*), then *i* will not form a bond in this time step, even if it has other possible bond partners. This crosslinking phase is then followed by a final cool-down, from 100 K to 0 K during another 3×10^5^ timesteps, with no new crosslinking. To mimic the tensioned structure of hydrated gels, the simulated polymer network is again geometrically transformed (in the manner described above) prior to introducing swimming cells: to impose mechanical tension, the coordinates are uniformly expanded by a factor of 1.2. This procedure yields a starting conformation that is both dense and under tension, reflecting the structural characteristics of real hydrogels, which swell under hydration. Subsequently, the system is allowed to mechanically equilibrate for an additional 3×10^5^ time steps at 0 K.

### Simulation procedure

Within the constructed gel matrix, a swimming cell is introduced at a random location, ensuring no overlap with the gel matrix by briefly relaxing any intersecting gel beads. To mimic bacterial swimming, including both rotational and translational motion, a torque of 500 pN.nm about its axis and an external force of 0.1 pN along its axis are applied, corresponding to approximately one full rotation per cell-length of forward displacement during free swimming. For the simulation of varied curvatures, a smaller torque (100 pN.nm) is applied. The motion of the swimming cell in the gel is then simulated at 0 K, and the positions and velocities of all cell beads are recorded every 500 timesteps (each timestep is 0.5 µs) for subsequent analysis. Each simulation is ran for 2×10^7^ timesteps. To determine when a swimming cell stopped moving, displacements are calculated over a time interval of 10^4^ timesteps. At each frame, the Euclidean distance between the current position and the position 10^4^ timesteps later is computed. If this displacement is less than 2 µm for 4×10^4^ consecutive timesteps, the cell is considered to have stopped. The first frame meeting this condition is recorded as the stopping point. If no such event occurs, the cell is classified as motile throughout the trajectory. A total of 1000 simulations per condition (e.g. per curvature) is done.

### Gel shear modulus measurement

Following an initial equilibration phase and a relaxation phase of the gel as described above, shear deformation is introduced by applying an affine box deformation in the *xy* plane at a constant strain rate of 0.001 ms^−1^. During the subsequent simulation, which lasts 5×10^5^ time steps at 0 K, the shear stress *σ_xy_* of the gel matrix, box tilt *L_xy_*, and box length *L_y_* (see SI Figure 6c) are recorded every 500 time steps. The shear strain is calculated as *γ* = *L*_*xy*_⁄*L*_*y*_, and the shear modulus is determined from *G* = Δσ_*xy*_ ⁄Δ*γ*_*xy*_.

### Cell curvature measurement

To estimate cell curvature, we fit a circle that passes through two central body atoms and is tangent to the directions of the two arms (see SI Figure 6b). Specifically, we first extract two short segments representing the left and right arms of the cell, each comprising 3–4 beads. Linear regression is used to fit a line to each arm segment, yielding estimates of their orientations. Two beads located near the midpoint of the body — one at the junction with each arm — are used to define the arc of the cell body. We then compute the position and radius of the circle that passes through these two central atoms and is tangent to the previously fitted arm directions. The circle is determined by minimizing a cost function that penalizes deviation from both distance constraints (points lying on the circle) and angular constraints (tangency with arm directions), using nonlinear optimization. The fitted radius is used to quantify cell curvature.

## ACKNOWLEDGEMENTS

The 3D-tracking MATLAB code was graciously provided by Katja M. Taute. This research project was funded by the START-Program of the Faculty of Medicine of the RWTH Aachen University and by the Deutsche Forschungsgemeinschaft (DFG, German Research Foundation) – Project 570781731. This research was also supported by the National Institute of General Medical Sciences (NIGMS) of the National Institutes of Health (NIH) under Award Number R01 GM082938 (NSW). The content is solely the responsibility of the authors and does not necessarily represent the official views of the National Institutes of Health. This work was supported in part by the National Science Foundation, through the Center for the Physics of Biological Function (PHY-1734030).

## DATA AVAILABILITY

All the bacterial trajectories from the main chemotactic experiments will be made publicly available on the repository RWTHPublications, as well as swim extent data (both experimental and in silico) and supplementary movies, upon publication of this article.

