## Supplementary Information for "Cell-body curvature modulates stall frequency to enhance *Vibrio cholerae* swimming and chemotaxis through hydrogels"

**Supplementary Discussion: on the choice of PVP K90 and soft agar hydrogels as simple mucus-mimicking media.**

Mucus thickness, composition, and viscoelastic properties differ widely, from the thin layer protecting our eyes to the several hundred-micron thick multilayered lining in our colon[19–22]. Mucus structure at the bacterial scale ranges from an almost purely viscous solution to a highly confining hydrogel network with sub-micron pore sizes[23] and scale-dependent rheological properties[21,24,25]. The mucus of interest here is the loosely attached small-intestine mucus, as it is the barrier that *V. cholerae* must cross to reach its infection sites deep into small-intestine crypts[2]. Small-intestine mucus contains gelling mucins MUC2 and displays a mesh structure. In the healthy human small intestine, shear moduli were recently reported just under 10 Pa[27], in the lower range of the typical 1-100 Pa span reported along the whole intestine in various animal models[26]. Recent estimates of median pore sizes of canine small-intestinal mucus (using Cryo-SEM) are close to 2-3.5 µm. The bulk viscosity of human or pig small-intestine mucus is within 60-5,000 cP[25], with shear-thinning properties.

Agarose hydrogels display a nonlinear relationship between percentage and average pore size, with a widening distribution at lower percentages (e.g. see Pernodet et al.[28] Fig. 5). The exact porosity of agar, a lower purity polymer of agarose of ill-characterized molecular weight (which can influence pore size further[29]), is therefore hard to predict. Estimates are in the range of pores close to bacterial size; Licata et al.[12] estimated an average pore size for 0.25% agar in the 0.7-4.8 µm range. The elastic component (shear modulus) of 0.2% agar is expected to be close to that of 0.2% agarose (recently measured at 260 Pa[14]).

Solutions of polyvinyl pyrrolidone (PVP), a linear unbranched polymer with different molecular weights available, are widely used to study the relationship between viscosity and bacterial motility[9,10,30], but do not recapitulate the mesh structure of intestinal mucus. PVP K90 (average molecular weight of 360 kDa) has been demonstrated to display non-Newtonian shear thinning rheological properties down to microbial motility scale[10]. Solutions up to 40 cP can be easily and repeatability handled in the laboratory.

We therefore used here mostly 0.2% agar hydrogels, as well as PVP K90, as two rheological extremes of mucus-mimicking media. PVP K90 lacks the hydrogel structure but may reflect the viscous non-Newtonian non-gelling (very low shear modulus) mucus while agar hydrogels may reflect the hydrogel mesh structure but with a higher shear modulus (lower deformability) than at-rest small-intestine mucus. We then leveraged an in-silico model to extend our experimental agar results to mesh with realistic small-intestine shear moduli.

| strain | Total number of motile trajecto-ries | # of analyzed trajecto-ries | Cumul. duration (s) | Motile speed (µm/s) | Turning freq. ($s^{-1}$) | Forward duration (s) | Backward duration (s) |
|---|---|---|---|---|---|---|---|
| wt | 6,897 | 3,554 | 9,547 | 84.8 ±1.6 | 1.30 ±0.13 (12,921) | 0.46 ±0.03 (2,810) | 0.156 ±0.002 (3,790) |
| Δ*crvAB* | 5,305 | 2,796 | 7,999 | 81.5 ±1.0 | 1.32 ±0.06 (10,708) | 0.47 ±0.02 (3,336) | 0.156 ±0.007 (4,500) |

SI Table 1 – **Detailed dataset of wt and Δ*crvAB* strains' random swimming motility across three replicates in buffer.** Displayed values are cumulative sum (total number of trajectories and analyzed trajectories) or averages ± standard deviations across independent replicates. The total numbers of events across replicates are indicated in parentheses. Bacteria are deemed motile if their average swimming speed exceeds 15 µm/s. Trajectories are analyzed when exceeding 0.8 s in duration and 15 µm/s in average swimming speed. Forward and backward runs were identified by their bordering turning angles (see Methods). The detailed data here corresponds to analysis results displayed in Figure 1.

| Conditions | Strain | Number of independent replicates | Motility threshold (µm/s) | Chemotactic drift $v_d$ (µm/s) | Number (duration) of motile trajectories |
|---|---|---|---|---|---|
| Buffer | wt | n=10 | 15 | 17.8 ±.9SE | 48,894 (51,710 s) |
| | Δ*crvAB* | | | 18.1 ±.7SE | 48,062 (53,822 s) |
| Agar .19-.21% | wt | n=7 | 0 | 0.45 ±.08SE | 30,805 (272,662 s) |
| | Δ*crvAB* | | | 0.26 ±.05SE | 21,069 (216,381 s) |
| PVP K90 5% (w/v) ~38 cP | wt | n=5 | 4 | 1.70 ±.11SE | 5,101 (31,754 s) |
| | Δ*crvAB* | | | 1.81 ±.17SE | 4,108 (25,589 s) |

SI Table 2 – **Detailed chemotaxis of the main datasets of wt and Δ*crvAB* strains in 200 µM/mm serine gradients.** These datasets are at the root of analysis displayed in Figure 2 and SI Fig. 2ab (buffer), Figure 2 and 3 (agar), Figure 2 and SI Fig. 2c-d (PVP).

| Conditions | Shape | # of swim phases | Exponential decay length estimate $\lambda$ (µm) | Ratio of exponential lengths $\lambda_{curved}/\lambda_{straight}$ |
|---|---|---|---|---|
| Experimental (agar ~.20%) | Curved (wt) | 70,545 | 28 ±1.7 (SD, n=7) | 1.26 ±0.10 (SD, n=7) |
| | Straight (Δ*crvAB*) | 60,727 | 22 ±1.7 (SD, n=7) | |
| In silico (~100 Pa) | Curved | 1000 | 100 ±6 (95%CI) | 1.25 |
| | Straight | 1000 | 80 ±5 (95%CI) | |

SI Table 3 – **Summary of exponential decay length estimates from experimental and in-silico survival distributions of swimming phase extents.** In experimental data, the decay length is estimated for each phenotype on each experimental replicate and the noise reflects the standard deviation over the n=7 replicates. In in-silico data, the slope is estimated for each shape over its full distribution (for n=1000 simulations) and the noise reflects the 95% confidence intervals. The estimates and their 95% CI were obtained using the expfit function (MATLAB R2023a). The in-silico parameters were curvatures of 0 and 0.71 $\mu m^{-1}$ for the straight and curved cell shapes respectively, in a mesh with a shear modulus of 95.2 Pa. The detailed data here corresponds to analysis results displayed in Figure 4a.

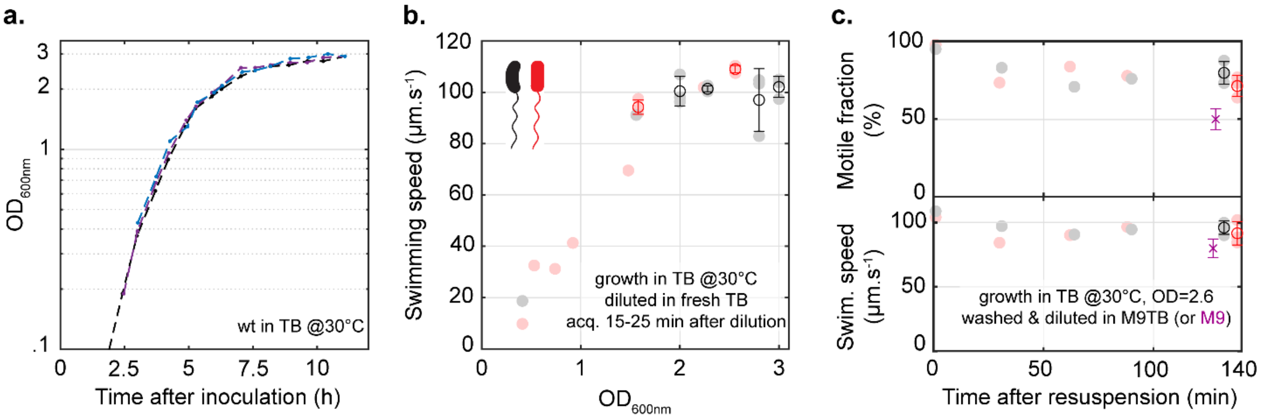


SI Figure 1 – **Growth and acquisition conditions ensuring stable motility parameters over acquisition times. (a)** Growth curve of wt strain in TB at 30°C 250 rpm, measured via optical density at 600nm ($OD_{600nm}$) over three independent experiments. **(b)** Swimming speed of motile fraction as a function of $OD_{600nm}$. The culture is diluted to an approximate $OD_{600nm}$=0.005 in fresh TB, then a 3D-tracking acquisition is made after 15-25 min to allow for adaptation. Points represent technical replicates over a single experiment. **(c)** Motile fraction and swimming speed over time, in final growth and acquisition conditions chosen (except for $OD_{600nm}$=2.6 instead of 2.2-2.4). The motility parameters are stable beyond the maximum duration after dilution accepted in further experiments (120 minutes). Points represent technical replicates over a single experiment. M9 salts + 0.01% TB (M9TB) was chosen as motility medium instead of M9 salts alone (purple point, wt strain) due to a clear decrease in motile fraction in the latter.

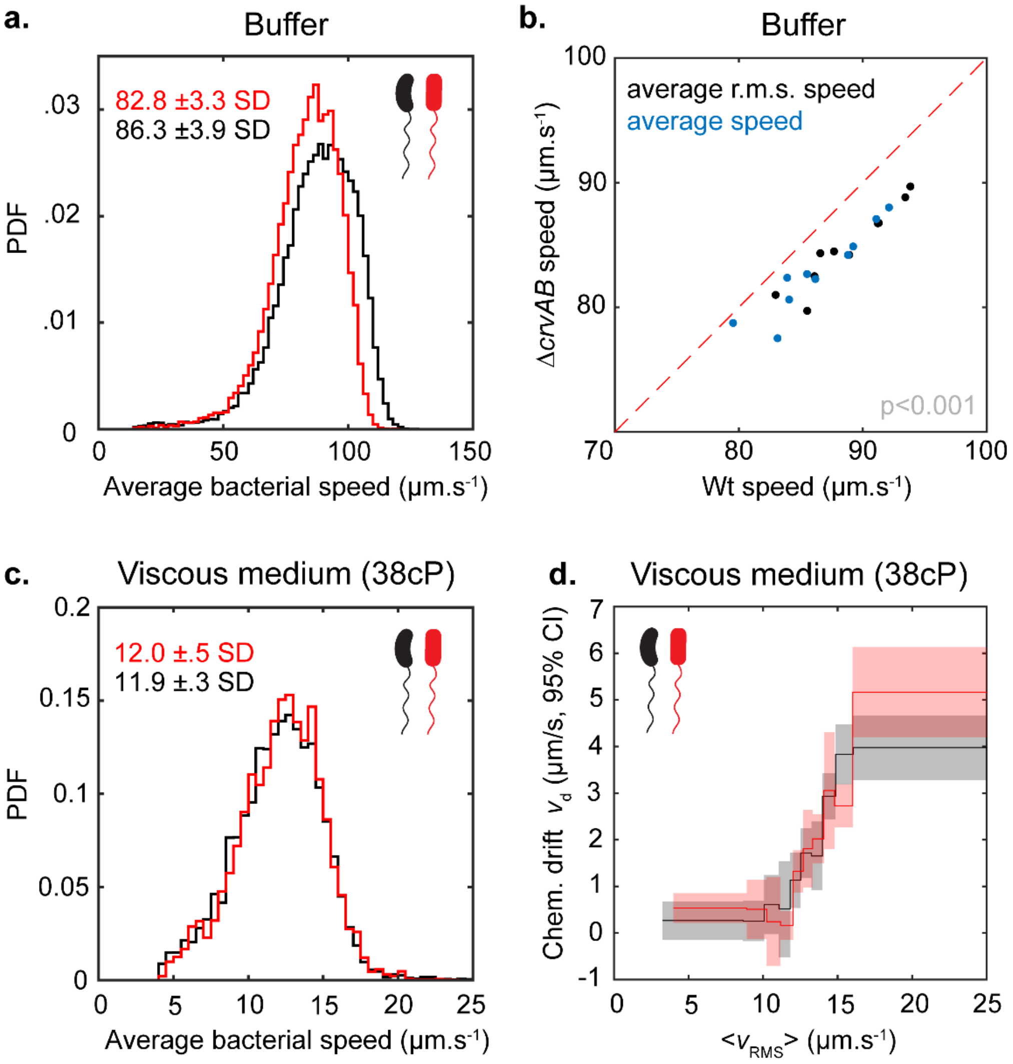


SI Figure 2 – **Further navigation properties in chemotaxis experiments in simple buffer or in PVP K90 5% (~38 cP).**

**(a)** Average bacterial swimming speed distributions and averages and standard deviations over n=10 replicates.

**(b)** Average swimming speed and root-mean-squared (r.m.s.) swimming speed of wt against Δ*crvAB* showing a small but repeatable increase of swimming speed for the curved-cell shape relative to the straight-cell one (on average 4.2 (±0.6 95% CI)%).

**(c)** Average bacterial swimming speed distributions.

**(d)** Drift velocity per RMS swimming speed bin, each bin determined to contain 10% of the total population.

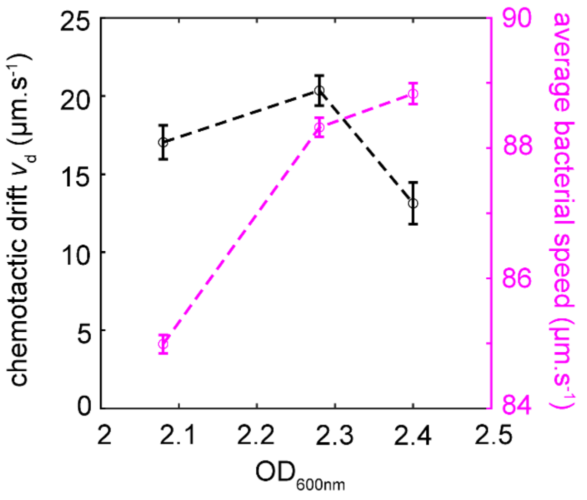


SI Figure 3 – **Chemotactic drift variability in buffer.** Chemotactic drift (black, left) and average swimming speed (magenta, right) for bacterial populations from one wt bacterial culture probed over three different OD values, following the same protocol as for regular chemotaxis experiment in buffer. Error bars represent the standard deviation between three successive technical replicates. While OD=2.2-2.4 was selected to ensure stable and high motile fractions (see SI Figure 1), chemotactic drifts showed a decrease as the OD increased. This imposed our strict requirement on identical OD (±0.01) within the same experiment to compare the strains (see Methods) and probably underlines the observed dispersed performances between experiments (Figure 2b).

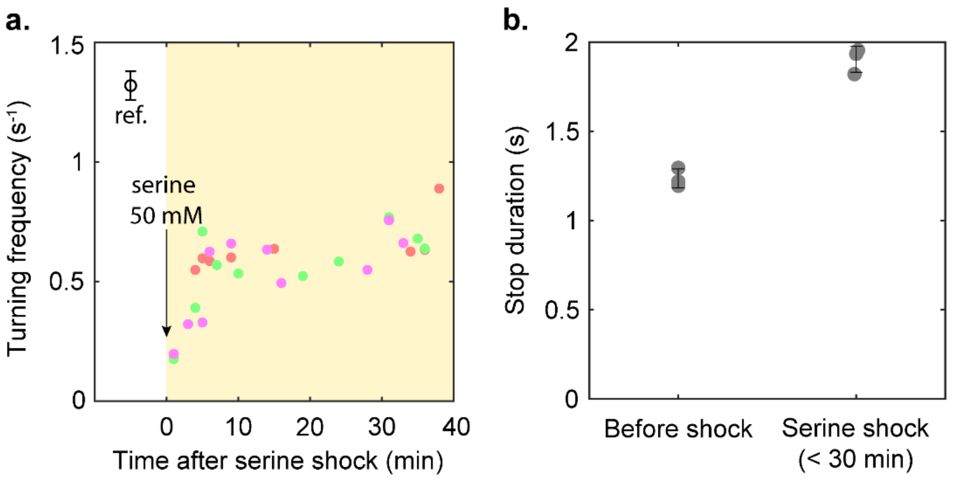


SI Figure 4 – **Data supporting turning as the primary mechanism for escaping stalls in hydrogels. (a)** Turning frequency of wt strain in M9TB after sudden exposure to 50 mM L-serine (so-called serine shock). **(b)** Stop durations of wt in soft agar in a chemotactic chamber under typical assay conditions (200 µM/mm serine gradient in M9TB, “before shock”), then after replacement of the reservoirs’ content with M9TB + 50mM serine (“serine shock”, within 30 min).

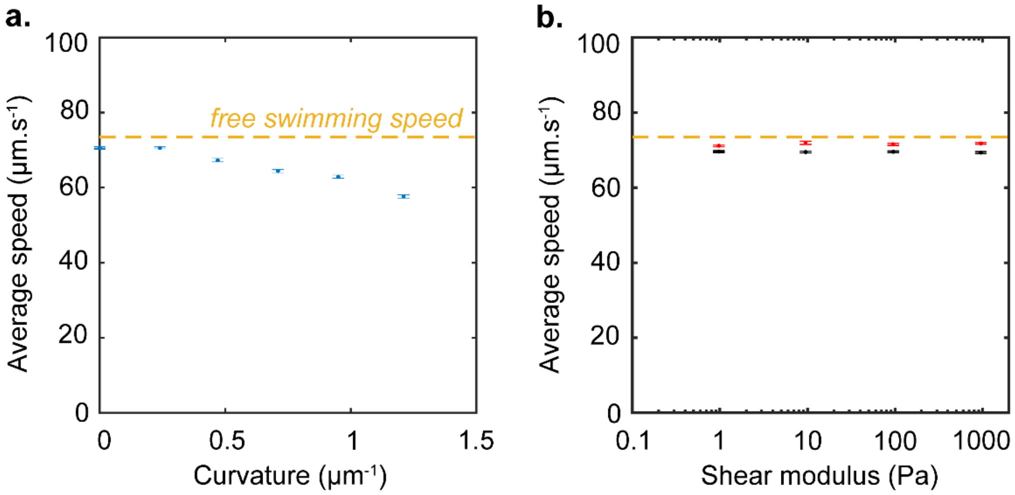


SI Figure 5 – **Average swimming speeds of "bacteria" in in-silico coarse-grained molecular dynamics simulations.** The average speeds and their 95% confidence intervals are represented over either **(a)** the curvature range tested or **(b)** over the range of stiffnesses tested for both straight (red) and curved (black) "bacteria".

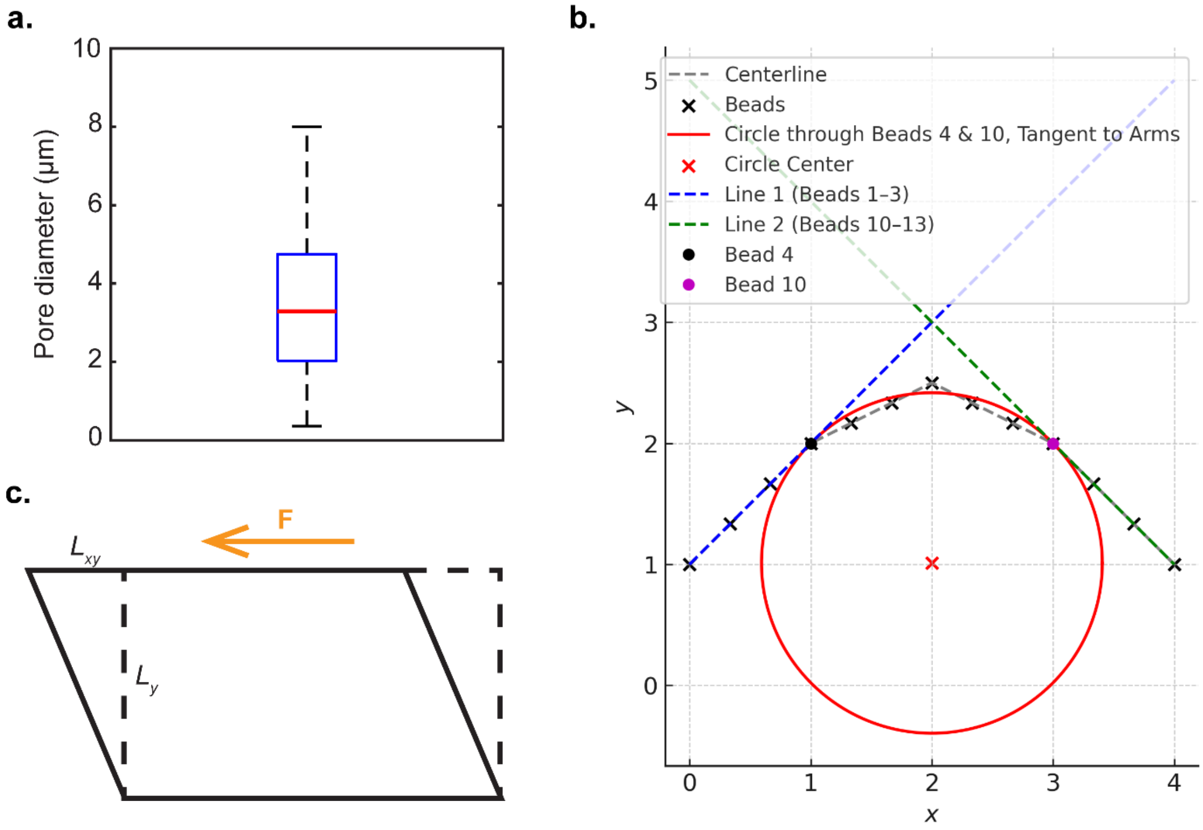


SI Figure 6 **– Details of parameters in our in-silico coarse-grained molecular dynamics model. (a)** Pore size distribution with an average pore size of 3.49 μm (±1.79 μm SD). This falls within estimated range of mean pore size for soft (0.25%) agarose hydrogels of 0.7-4.8 μm and close to recent estimates of median pore sizes of canine small-intestinal mucus (using Cryo-SEM) of 2-3.5 μm. **(b)** Estimating in-silico bacteria curvature (complementary illustration for Methods). **(c)** Measurement of in-silico mesh's shear modulus (complementary illustration for Methods).
